# Non-traumatic fMRI of identity-state expression and switching in co-conscious DID

**DOI:** 10.64898/2026.04.20.718646

**Authors:** Shogo Kajimura, Kenichiro Okano, Senkei Ueno, Junko Yamada, Haruna Fukui, Aiko Hirai, Ayahito Ito, Nobuhito Abe, Ryusuke Nakai, Shunichi Noma, Toshiya Murai

## Abstract

Dissociative identity disorder (DID) remains debated because identity-state phenomena are privately experienced and may be attributed to suggestion, simulation or role enactment. Most neuroimaging studies rely on symptom provocation or traumatic recall, which complicates interpretation and is poorly suited to co-conscious presentations where simultaneous awareness should make state differences hardest to detect. We applied Identity-State Characterization and Analysis (ID-SCAN), a non-traumatic, identity-cued functional magnetic resonance imaging protocol, to a DSM-5-diagnosed woman with persistent co-consciousness between Adult and Adolescent identity-states. One task used identical insect images that evoked opposite preferences across identity-states; the other used trait judgments about self, the other identity-state and a shared intimate other. Analyses combined Bayesian single-case general linear modelling, generalized psychophysiological interaction connectivity and searchlight representational similarity analysis. Identity-instruction cue epochs were pooled across tasks to assess switch direction. The same insect stimuli engaged different valuation-related configurations across identity-states: Adolescent-selective effects centred on striato-thalamic regions, whereas Adult-selective effects extended to amygdala and orbitofrontal/medial prefrontal cortex, with distinct task-evoked coupling. Adult-as-self and Adolescent-as-self occupied separable positions within canonical self-referential regions. Cue-locked activity differed by switch direction across tasks, with larger reconfiguration when switching to Adult (mean between-cluster beta separation 4.30 versus 0.85; permutation *p* = 0.0001). Cross-task overlap localized a limited shared task-related substrate mainly to posterior visual and dorsal parietal cortex. Even under persistent co-consciousness and without trauma provocation, identity-state expression and switching showed convergent within-person neural signatures. The findings support non-traumatic mechanistic phenotyping of dissociative presentations and motivate cohort and longitudinal studies, including treatment-tracking work.

## Introduction

Dissociative identity disorder (DID) remains among the most debated diagnoses in psychiatry Its defining phenomena are experienced privately and can be dismissed as suggestion, simulation or iatrogenic role enactment.^1–3^ Within a single nervous system, shifts in who one takes oneself to be, what one feels and what one wants or fears can be experienced as abrupt and pervasive, yet outsiders have access only to testimony and observable behaviour. This epistemic asymmetry has kept DID at the margins of mechanistic psychiatry and has discouraged the use of dissociative self-states as a model for studying how conscious selfhood is implemented in brain networks. At the same time, DID is associated with severe functional impairment, self-harm and high treatment utilisation,^4,5^ patterns that are difficult to explain if the disorder were primarily a product of deliberate performance. If DID reflects a genuine disturbance of conscious self-organization, its identity-state shifts constitute an unusually strong perturbation of subjective experience that could be used to link phenomenology to reproducible neural mechanisms. Progress therefore requires paradigms that move beyond testimonial validity and treat identity-states as quantifiable, reproducible configurations within a single brain.

Functional neuroimaging studies have begun to address this challenge by showing identity-dependent activation and connectivity profiles in limbic, paralimbic and default-mode systems,^7–9^ and by demonstrating that these profiles are not reproduced by healthy controls instructed to simulate DID.^9,10^ The theory of structural dissociation conceptualizes DID as an organization of personality into parts that differ in affective and defensive tendencies and that coexist with partially overlapping access to memory and agency,^6,44^ predicting that identity-states should be implemented as distinct, recurrent network configurations rather than as transient fluctuations of a unitary self. However, much of the existing literature relies on symptom provocation or traumatic autobiographical material, which raises ethical and interpretive difficulties: traumatic memories may be encoded and retrieved via mechanisms that differ from ordinary recollection,^15–17^ and the associated distress can increase susceptibility to participant expectancy and task-demand effects. There is therefore a practical need for tasks that are non-traumatic, standardizable and tightly linked to neural systems that can be interrogated in a single individual.

Co-consciousness, defined as simultaneous awareness by more than one identity-state, is recognized in clinical descriptions and treatment guidelines^11,12^ but further complicates neurobiological interpretation. Most imaging studies have focused on patients whose switches are accompanied by marked inter-identity amnesia, leaving open the question of how identity-states are instantiated when co-consciousness is the typical mode of experience.^7–10,13,14^ Under co-aware conditions, sustained blending between identities should attenuate contrasts, and the phenomenology can resemble deliberate role play because both involve monitoring which state is being expressed and how one is expected to respond. These features motivate tasks that are difficult to simulate strategically on a trial-by-trial basis and that can generate within-person replications across domains. It is equally important to characterize what is preserved across identities so that identity-selective effects can be interpreted against an explicit task-general baseline rather than attributed to global differences in attention, effort or compliance.

Here we present an in-depth single-case fMRI study of a woman with DID and persistent co-consciousness between an Adult and an Adolescent identity-state. The participant reported no inter-identity amnesia in ordinary activities, and the Adolescent was typically foregrounded in daily life; this presentation provides a stringent test case because co-awareness should, if anything, attenuate identity-state differences. We developed an Identity-State Characterization and Analysis (ID-SCAN) protocol that triangulates identity-state representations and switching dynamics across two non-traumatic, identity-cued judgment tasks and cross-task analyses. In Experiment 1, we used naturally occurring Adult-Adolescent differences in affective preferences to probe identity-dependent valuation and salience processing with identical insect stimuli.^18,19^ In Experiment 2, we adapted a trait-judgment paradigm to examine how each identity represents self, the other identity and an intimate other within canonical self-referential regions, using representational similarity analysis (RSA) and multidimensional scaling (MDS) to map identity-dependent person-knowledge geometry.^20–23^ Because self and close-other representations in this network show reliable structure that tracks social connection and preserves self-specificity,^20–22^ this framework provides an interpretable test of whether each identity-state is instantiated as a distinct self-referential representation rather than a generic response style. We also used the commonality of task components (both tasks were identity-cued and required explicit judgments) to conduct a pooled analysis of the identity-instruction epoch to quantify task-general switching dynamics, and a cross-task overlap analysis to identify a conservative within-person baseline that constrains interpretation of identity-selective effects.

## Materials and methods

### Participant, ethics and consent

A right-handed woman with DID according to DSM-5 criteria participated.^28^ Two identity-states were examined: an Adult identity-state (Adult; denoted A in figures) and an Adolescent identity-state (Adolescent; denoted Y). The participant reported persistent co-consciousness with no inter-identity amnesia in daily life; the Adolescent was usually foregrounded, whereas the Adult was experienced as more fatigued and less active. At scanning, she had comorbid major depressive disorder and was taking duloxetine (20 mg). The protocol was approved by the Kyoto Institute of Technology ethics committee (approval No. 2024-06). All procedures contributing to this work comply with the ethical standards of the relevant national and institutional committees on human experimentation and with the Helsinki Declaration of 1975, as revised in 2013.^29^ Written informed consent was obtained from the participant, who consented to the inclusion of anonymized material relating to herself. A safety plan was implemented for all visits; during the first session, the participant’s attending physician was present on site to confirm clinical stability and support predefined stopping criteria.

### MRI acquisition and preprocessing

Task fMRI data were acquired on a 3.0-T Siemens Prisma with a 32-channel head coil using the Brain/MINDS Beyond Harmonized MRI Protocol (HARP).^30^ The functional sequence was a gradient-echo EPI (TR = 0.8 s, TE = 34.4 ms, flip angle = 52 degrees, field of view = 206 x 206 x 144 mm, matrix size = 86 x 86, 60 axial slices, slice thickness = 2.4 mm, no gap, multiband factor = 6). A T1-weighted MPRAGE structural scan was acquired with TR = 2.5 s, TE = 2.07 ms, flip angle = 8 degrees, and 0.8-mm isotropic voxels. To correct susceptibility-induced EPI distortions, a dual-echo gradient-echo field map was acquired (Siemens gre_field_mapping), yielding two magnitude images (TE1 = 5.39 ms; TE2 = 7.85 ms) and a phase-difference image (delta-TE = 2.46 ms; TR = 0.336 s; flip angle = 60 degrees; slice thickness = 2.4 mm; no gap).

Preprocessing was conducted in SPM12^31^ and included slice-timing correction (reference: median acquisition-time slice), realignment, field map-based unwarping, coregistration to the T1 image, normalization to MNI space via unified segmentation, and resampling to 2-mm isotropic voxels. Field map phase-difference and magnitude images were processed using SPM12’s FieldMap toolbox to derive voxel displacement maps, which were applied during the unwarping step. No spatial smoothing was applied; the Bayesian estimation framework used for all activation analyses estimates the spatial noise structure via an internal spatial prior (uninformative global Laplacian, UGL), making conventional Gaussian smoothing unnecessary and potentially detrimental to the Bayesian posterior probability maps. A 128-s high-pass filter was applied, and six motion parameters per run were included as nuisance regressors. No volume censoring (scrubbing) was performed. Framewise displacement (FD) was computed using the Power *et al*. formula (50-mm head radius). Across the six task runs contributing to the analyses (three for Experiment 1, three for Experiment 2), mean FD was 0.28 mm for Experiment 1 (range across runs: 0.27-0.30; maximum single-volume FD: 6.68 mm; 7.7% of volumes exceeding 0.5 mm) and 0.42 mm for Experiment 2 (range: 0.39-0.43; maximum: 2.50 mm; 27.3% exceeding 0.5 mm). The higher motion in Experiment 2 was driven by sustained low-amplitude drift rather than spike artefacts; the maximum single-volume FD in Experiment 2 (2.50 mm) was lower than in Experiment 1 (6.68 mm).

### Experiment 1: identity-cued preference/aversion task

Experiment 1 targeted the valuation component of the ID-SCAN framework and addressed whether identity-specific neural patterns are robust across tasks and difficult to mimic by deliberate simulation. We selected insect stimuli because interviews confirmed a marked and stable Adult-Adolescent divergence in attitude (aversion versus liking), enabling rapid preference appraisals with benign content while holding stimulus input constant across identities. This design feature makes it difficult to reproduce identity-consistent responding strategically on a trial-by-trial basis.

#### Stimuli selection and post-scan rating

Semi-structured interviews identified three insect categories (spiders, Cephonodes hylas, and dragonflies) for which Adult reported dislike, whereas Adolescent reported liking (Fig. 1B, C). For each category, seven photographs were obtained from the iNaturalist database^32^ and paired with phase-preserving grid-scrambled versions, yielding 21 insect and 21 scrambled images. After scanning, each image was rated on a visual-analog scale (0-100) separately in each identity-state. As a manipulation check, we compared mean liking ratings between Adult and Adolescent across images (Fig. 1D). We additionally verified that the Adolescent-Adult liking difference was positive for every insect image (i.e. no insect image received a higher rating in Adult than in Adolescent).

**Figure 1.**
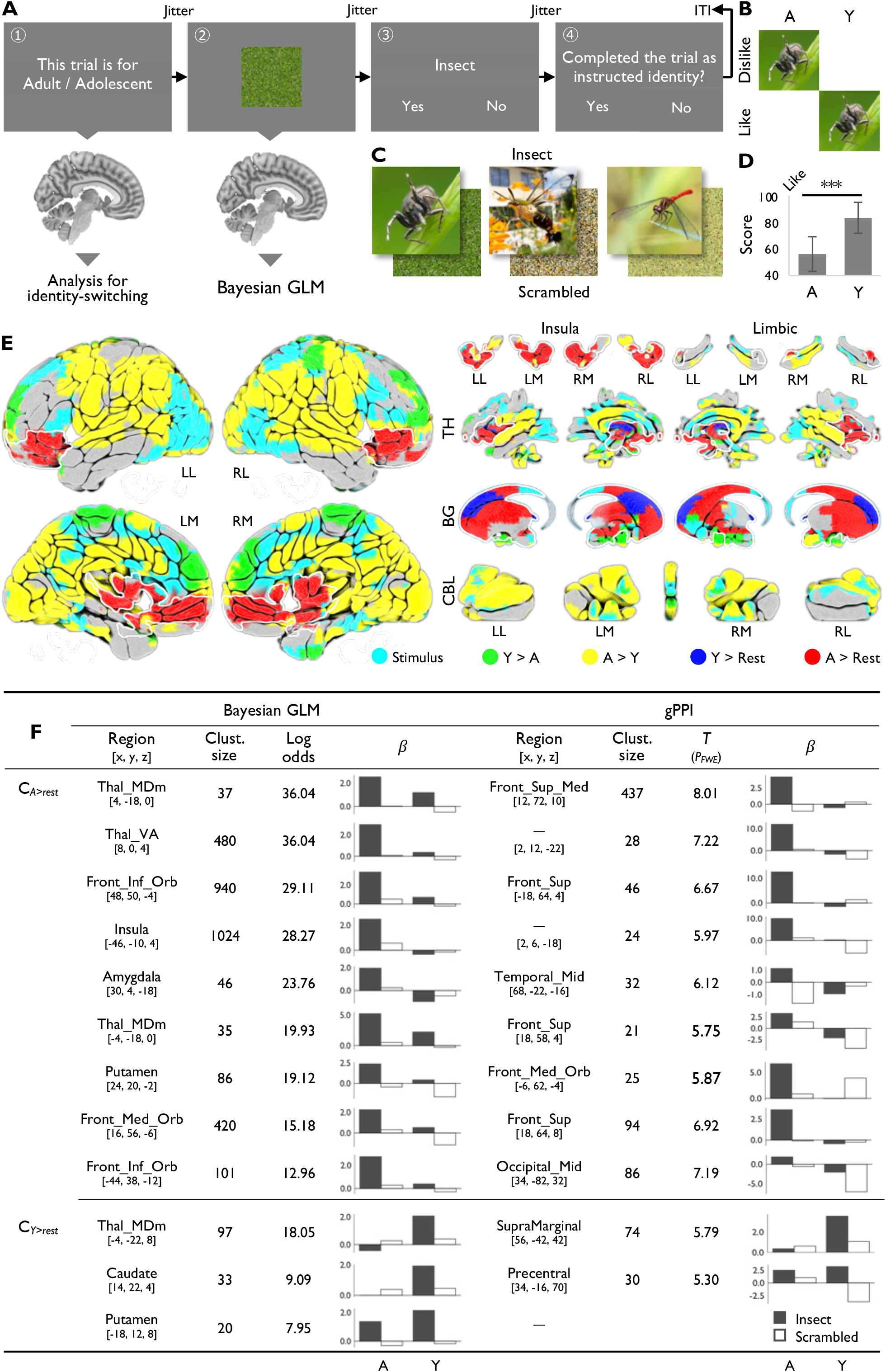
Methods and results of Experiment 1. (A) Schematic of a single trial in the identity-cued preference/aversion task. At the start of each trial, an on-screen cue instructed which identity-state to foreground; this cue period constitutes the identity-instruction epoch pooled across tasks in the task-general switching analysis. The following image-presentation epoch is the segment modelled in the Bayesian first-level GLM for Experiment 1. (B) Summary of the relationship between identity-state and affective attitude toward the stimulus categories used in Experiment 1. (C) Example stimuli. Representative intact insect images and their phase-preserving grid-scrambled counterparts for spiders, Cephonodes hylas and dragonflies. (D) Post-scan preference ratings. Error bars indicate within-stimulus variability, and the asterisk denotes a significant difference between Adult and Adolescent ratings (paired *t*-test across stimuli). (E) Surface rendering of the results of Bayesian GLM contrasts. PPM thresholding: identity-selective spotlight contrasts were thresholded at log-odds >= 4.6 within the a priori valuation mask with extent >= 20 voxels; all other Bayesian GLM maps were thresholded at log-odds >= 10 with extent >= 20 voxels. The white outline indicates the a priori affect/valuation mask (AAL3) within which these spotlight contrasts were evaluated, encompassing bilateral orbitofrontal, insular, striatal, thalamic and midbrain regions. (F) Summary of insect-selective valuation clusters and gPPI connectivity seeds. Adu: Adult, Ado: Adolescent, ITI: inter-trial interval, LL: left-lateral view, RL: right-lateral view, LM: left-medial view, RM: right-medial view, TH: thalamus, BG: basal ganglia, CBL: cerebellum. Regional labels in (F) follow AAL3 nomenclature, including Thal_MDm (mediodorsal nucleus of the thalamus), Thal_VA (ventral anterior nucleus of the thalamus), Front_Inf_Orb (inferior frontal gyrus, orbital part), Front_Med_Orb (medial orbitofrontal cortex), Front_Sup_Med (medial superior frontal gyrus) and Occipital_Mid (middle occipital gyrus).

#### Trial structure

During fMRI, each trial consisted of (i) a 3.0-s on-screen identity cue (‘Adult’ or ‘Adolescent’) followed by a jittered fixation (uniform, 2.0-3.0 s), (ii) 2.0-s picture presentation (insect or scrambled) followed by a second jitter (2.0-3.0 s), (iii) a forced-choice perceptual judgment (‘insect or scrambled?’; 5.0-s response window) followed by a third jitter (2.0-3.0 s), and (iv) an identity-check screen (‘Did you perform this trial in the instructed identity-state?’; yes/no; 5.0-s window) followed by a jittered inter-trial interval (3.0-5.0 s).

#### Identity-cued Bayesian GLM and spotlight contrasts

First-level analyses used SPM12’s Bayesian GLM with four regressors per run corresponding to the 2 x 2 design (Adult/Adolescent x insect/scrambled), yielding 14 insect and 14 scrambled trials per identity-state per run (84 insect and 84 scrambled trials in total across three runs). Events were modelled at picture onset with the canonical haemodynamic response function (no temporal or dispersion derivatives); motion parameters and session constants were included as nuisance regressors. Temporal autocorrelation was modelled with an AR(3) process during Bayesian estimation, with uninformative global Laplacian (UGL) priors for both signal and noise. We tested main effects and identity-selective ‘spotlight’ contrasts that isolated insect-selective responses within each identity (weights: Adult-insect [3, −1, −1, −1]; Adolescent-insect [−1, −1, 3, −1]). Spotlight maps were evaluated within an a priori valuation mask (AAL;^33^) spanning orbitofrontal, insular, striatal, thalamic and midbrain regions. Within this mask, spotlight posterior probability maps were thresholded at log-odds >= 4.6 (posterior probability >= 0.99) with an effect-size threshold of 0.1% signal change and cluster extent >= 20 voxels. The lower log-odds threshold for the a priori masked spotlight contrasts, relative to the whole-brain maps, follows the logic of small-volume correction: because the search space is restricted to a theoretically motivated region, a less stringent threshold maintains sensitivity while controlling false-positive rates within the mask. All other Bayesian GLM maps used log-odds >= 10 (posterior probability >= 0.99995) with the same effect-size and extent thresholds.

#### Identity-dependent connectivity

To examine whether spotlight activations were embedded in different functional networks, generalized psychophysiological interaction (gPPI) analyses were conducted using the SPM12 PPPI toolbox.^34^ Seeds were defined as first eigenvariates extracted from local-maximum clusters identified by the spotlight contrasts. For each seed, the gPPI model included the seed time series (physiological regressor), four psychological regressors (Adult-insect, Adult-scrambled, Adolescent-insect, Adolescent-scrambled), and one interaction regressor per condition. Motion parameters and a 128-s high-pass filter matched the activation model. Identity-selective PPI contrasts mirrored the activation spotlight contrasts. Seed-wise results were assessed at voxel-wise *p* < 0.001 (uncorrected) with cluster-wise family-wise error correction *p* < 0.05 (cluster extent > 20 voxels). This frequentist thresholding was adopted because the PPPI toolbox does not natively support Bayesian estimation; the combination of uncorrected voxel-wise and cluster-level FWE thresholds follows standard practice for single-subject gPPI analyses. Note that gPPI measures symmetric task-evoked functional coupling, not directed effective connectivity.

### Experiment 2: identity-cued trait-judgment task

Experiment 2 targeted person-knowledge representations within self-referential regions. Sixty personality adjectives were drawn from a Japanese Big Five adjective set.^35^ This trait-judgment paradigm exploits the well-described representational structure of canonical self-referential regions to test whether each identity instantiates a self-like representation, rather than expressing a generic response style under identity instruction.

#### Trial structure

During fMRI, each adjective trial consisted of (i) a 3.0-s identity cue (‘Adult’ or ‘Adolescent’) followed by a jittered fixation (2.0-3.0 s), (ii) three successive 3.0-s ratings (one per target), each separated by a 0.5-s blank screen and followed by a jittered inter-trial interval (3.0-5.0 s). The target label (‘Self’, ‘Other identity’ or ‘Intimate other’) was presented at the top of the screen, with an adjective in the centre and a 4-point Likert scale at the bottom (1 = strongly disagree, 2 = disagree, 3 = agree, 4 = strongly agree). Target order was randomized within trial. There were 120 trials (2 raters x 60 adjectives), yielding 360 rating events (~120 per run) across six conditions (Adult-rates-Adult, Adult-rates-Adolescent, Adult-rates-Other, Adolescent-rates-Adult, Adolescent-rates-Adolescent, Adolescent-rates-Other) in three runs.

#### Single-rating GLM

Single-rating beta images were estimated using a Bayesian GLM with one event regressor per rating (360 regressors in total across three runs, ~120 per run), convolved with the canonical HRF. Six motion parameters per run were included as nuisance regressors; AR(3) noise modelling, UGL priors and high-pass filtering matched Experiment 1.

#### Searchlight multi-regression RSA.^23^

Within 3-voxel-radius spherical searchlights (>= 8 voxels), we computed neural representational similarity matrices across all 360 ratings (Pearson correlation) and regressed the vectorized upper triangle of the neural similarity matrix onto vectorized binary model similarity matrices using ordinary least squares. Four model matrices were entered simultaneously: rater identity (alter versus host), target identity (alter, host, shared other), self-versus-non-self (self-judgments versus all other-judgments), and the full rater-by-target model. For statistical inference, we used 1,000 permutations per voxel (two-tailed): on each permutation, the rows and corresponding columns of the neural similarity matrix were shuffled to generate a null distribution of *t*-statistics under the null hypothesis of no systematic relationship between neural and model similarity structure; the permutation *P*-value was computed as the proportion of permutations for which the absolute permuted *t*-statistic equalled or exceeded the observed absolute *t*-statistic. Voxel-wise results were thresholded at *p* < 0.001 with cluster extent > 20 voxels (26-connectivity).

#### Visualization of person-knowledge geometry

To visualize representational geometry, we focused on the model that best explained similarity within the canonical self-referential processing network. We intersected its significance map with an independent meta-analytic mask from Neurosynth^36^ (search term: ‘judgments’; uniformity-test *z*-map thresholded at FDR *q* < 0.01) and computed a condition-level representational dissimilarity matrix for the six rater-by-target conditions. Metric MDS was applied to this 6 x 6 dissimilarity matrix. Dimensionality was set to three based on an elbow analysis of STRESS values (Fig. 2D).

**Figure 2.**
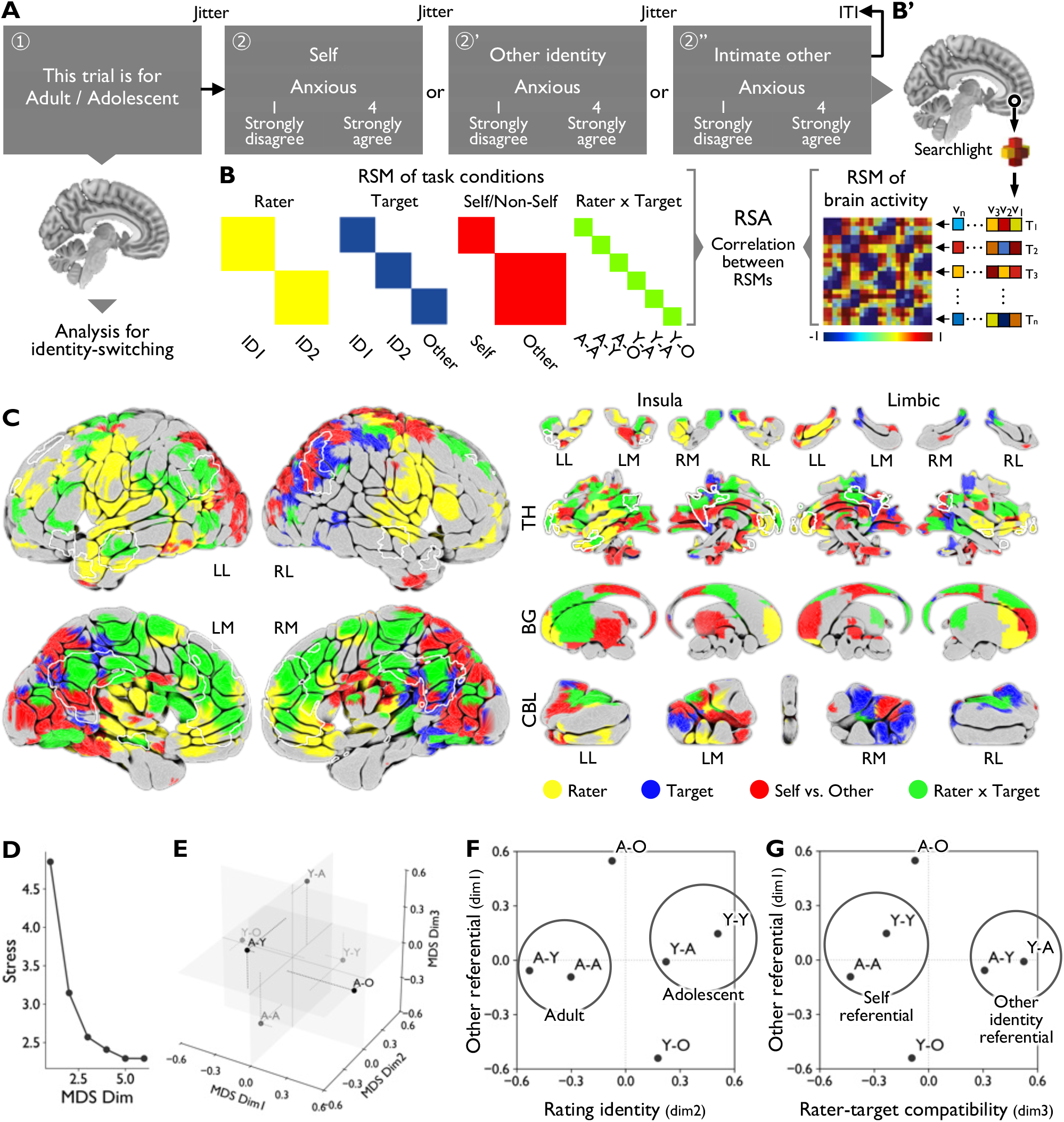
Methods and results of Experiment 2. (A) Schematic of a single trial in the identity-cued trait-judgment task. At the start of each trial, an on-screen cue instructed which identity-state (Adult or Adolescent) should serve as the rater; this cue period constitutes the identity-switching epoch pooled across tasks in the task-general switching analysis. The subsequent sequence of personality-trait ratings (Self, Other identity, Intimate other) is the segment modelled in the RSA and MDS analyses for Experiment 2. (B, B’) Task-structure-based representational similarity model and schematic of searchlight RSA. Neural RSMs were computed within spherical searchlights centred on each voxel and regressed onto this and other model RSMs. (C) Surface rendering of searchlight RSA results for the Rater x Target model. Voxels where the model significantly explained neural similarity structure (permutation-based voxel-wise *p* < 0.001, cluster extent >= 20 voxels) are shown; the white outline indicates the boundary of an independent meta-analytic ‘self-referential’ activation map from Neurosynth, which was used as an a priori mask for subsequent representational-geometry analyses. (D) Elbow plot of STRESS values as a function of MDS dimensionality (k = 1-6), showing a knee at k = 3, which motivated selection of a three-dimensional solution. (E) Three-dimensional MDS embedding of the Rater x Target conditions within the intersection of the Rater x Target RSA map and the self-referential mask. Each dot represents one condition (Adult-rates-Adult, Adolescent-rates-Adolescent, Adult-rates-Adolescent, Adolescent-rates-Adult, Adult-rates-Intimate other, Adolescent-rates-Intimate other), positioned according to a 6 x 6 condition-level representational dissimilarity matrix (correlation distance) derived from single-rating beta patterns in this ROI by averaging all rating pairs within each condition pair. (F) Two-dimensional projection of the same MDS solution onto dimensions 1 and 2. (G) Two-dimensional projection of the same MDS solution onto dimensions 1 and 3. ITI: inter-trial interval, A: Adult, Y: Adolescent, Adu-Adu: Adult-Adult, Adu-Ado: Adult-Adolescent, Adu-O: Adult-Other, Ado-Ado: Adolescent-Adolescent, Ado-Adu: Adolescent-Adult, Ado-O: Adolescent-Other, LL: left-lateral view, RL: right-lateral view, LM: left-medial view, RM: right-medial view, TH: thalamus, BG: basal ganglia, CBL: cerebellum.

### Task-general switching analysis

To isolate task-general signatures of intentional identity switching, we analysed the brief identity-instruction epoch at trial onset and pooled cue events across both tasks. Cue events were coded by the direction of switch (switch to Adult versus switch to Adolescent), independent of task content. In total, 203 cue events were modelled across six runs (84 from Experiment 1 and 119 from Experiment 2), comprising 101 switches to Adult and 102 switches to Adolescent. In Experiment 1, each trial included an identity-check screen (‘Did you perform this trial in the instructed identity-state?’; yes/no); the participant confirmed compliance on 96.4% of trials (81/84; three non-confirmations, all during Adolescent-cued trials). In Experiment 2, which did not include a forced-choice identity-check, all 119 trials were completed without reported compliance failures. Non-confirmed trials from Experiment 1 were coded as a separate error regressor. A Bayesian event-related GLM modelled cue epochs (3-s duration) using two regressors (switches to Adult, switches to Adolescent), a single regressor for error trials from Experiment 1, and motion parameters as nuisance regressors. AR(3) noise modelling and Bayesian estimation settings matched the preceding analyses. Primary contrasts tested task-general differences in cue-locked activity between switch directions, using thresholding matched to Experiment 1.

### Cross-task identity-invariant judgment analysis

To delineate an identity-invariant, judgment-related substrate shared across tasks, we combined identity-invariant maps derived independently from each experiment. In Experiment 1, the perceptual judgment period (Fig. 1A, phase 3) was modelled separately from picture viewing (Fig. 1A, phase 2) and a judgment > viewing contrast was computed for each identity-state; voxels reliably expressed in both states were summarized using a conjunction (Adult AND Adolescent) map thresholded as in the main Bayesian analysis. In Experiment 2, a rater-invariant target representation map was defined by inclusively masking voxels with significant Target identity and/or Self-versus-Non-Self effects while exclusively removing voxels with significant Rater identity and/or Raters x Targets effects (thresholded as in the main RSA). The Experiment 1 and Experiment 2 identity-invariant maps were intersected to identify convergent cross-task candidate regions supporting identity-invariant judgment.

## Results

### Experiment 1: preference/aversion judgment task

Experiment 1 tested whether the same benign stimuli would engage different valuation-related systems when the foreground identity changed. The participant confirmed performing 96.4% of trials in the instructed identity-state via the on-screen identity-check (81/84 trials; three non-confirmations, all during Adolescent-cued trials; these were modelled as error trials and excluded from activation contrasts). Post-scan ratings confirmed the intended manipulation: across insect stimuli, the Adolescent identity rated images as more likable than the Adult (paired *t*-test across images, *p* < 0.001). The insects versus scrambled contrast engaged visual cortex as expected, supporting data quality for identity-specific analyses (Fig. 1E). Collapsing across stimulus type, picture-viewing activity also differed between identity-states. Adolescent-cued viewing showed higher responses in dorsal and medial frontal cortex together with sensorimotor territories (Table S2), whereas Adult-cued viewing showed higher responses in orbitofrontal/medial prefrontal and striato-thalamo-cingulate and temporal systems (Table S3). These state-general differences provide an additional manipulation check that identity instruction induced broad reconfiguration during the viewing epoch, beyond stimulus-specific valuation effects.

Within the a priori valuation mask (Fig. 1E, white outline), identity-selective spotlight contrasts revealed dissociable valuation-related ensembles for the two identities when evaluating the same insect stimuli. Adolescent-selective insect responses were most prominent in striatal and thalamic loci (caudate/putamen and mediodorsal thalamus; Table S4). Adult-selective insect responses were broader, spanning right amygdala, orbitofrontal and medial orbital prefrontal cortex, insula, putamen and thalamic nuclei (Table S5).

Generalized psychophysiological interaction (gPPI) analyses showed that identity-selective spotlight nodes were embedded in dissociable task-evoked networks during evaluation of the same insect stimuli (Fig. 1F; Tables S4-S5). During Adolescent insect trials, striato-thalamic seeds identified by the spotlight contrast (caudate/putamen/mediodorsal thalamus) exhibited increased coupling with insula and medial temporal regions (e.g. parahippocampal cortex), together with action/attention-related territories including precentral and supramarginal cortex and dorsal frontal regions (Table S4). During Adult insect trials, seeds in amygdala and orbitofrontal/medial orbital prefrontal cortex, together with insula, putamen and thalamic nuclei, showed broad coupling with medial and lateral prefrontal cortex, dorsal parietal regions and temporo-occipital visual cortex (e.g. calcarine/cuneus/occipital/fusiform), as well as medial temporal structures (Table S5). Thus, identity-state differences were expressed not only in regional engagement within valuation circuitry but also in the pattern of task-evoked inter-regional coupling linking valuation hubs to perceptual and control systems.

### Experiment 2: trait-judgment task

Experiment 2 asked whether each identity would occupy a distinct self-like position within person-knowledge space, rather than merely producing a generic identity-cued response style. Multi-regression RSA demonstrated robust representational structure related to both rater identity and judged target (Fig. 2C). The rater identity model produced the broadest footprint, consistent with a widespread reconfiguration of person-knowledge representations when the foreground identity changed. Target-related structure was most prominent in posterior medial regions, including precuneus, with additional clusters extending into temporal and cerebellar regions. A Self-versus-Non-Self component was also detectable, indicating that self-related organization was preserved alongside rater-specific effects.

Within the canonical self-referential processing network (Fig. 2C, white line), the full rater-by-target model accounted best for representational similarity and defined the mask for geometric visualization. Metric MDS of the six rater-by-target conditions yielded a stable three-dimensional configuration that could be described by three interpretable dimensions (Fig. 2D-G). One dimension separated Adult-as-rater from Adolescent-as-rater across targets, consistent with a ‘rating identity’ axis. A second dimension separated self-judgments (Adult-as-self; Adolescent-as-self) from cross-identity judgments, consistent with ‘rater-target compatibility’, with judgments about the shared intimate other occupying an intermediate position. A third dimension captured perspective effects for the intimate other, consistent with an ‘other-referential’ axis. Thus, Adolescent-as-self occupied a distinct but self-like position rather than collapsing onto Adult-as-self or shifting toward other-judgments, indicating identity-specific self-representations within the self-referential system even under continuous co-consciousness.

### Task-general switching analysis

Cue-locked activity differed by direction of switching (Fig. 3B; Tables S6-S7). Switch-to-Adolescent > switch-to-Adult emphasized orbitofrontal/ventromedial prefrontal cortex, anterior insula and striatum, whereas switch-to-Adult > switch-to-Adolescent showed broader modulation including visual and temporal cortex, medial frontal/cingulate, insula, brainstem, cerebellum and sensorimotor regions. Cue-locked modulation was more extreme for switches to Adult: cluster-wise parameter estimates (median beta across voxels within each cluster, averaged across runs) separated Adult-cue beta values between Adult-ward versus Adolescent-ward clusters (mean difference = 4.30; permutation *p* < 0.0001) more than Adolescent-cue beta values (mean difference = 0.85; permutation *p* < 0.0001). Cluster boundaries were defined from the thresholded Bayesian contrast maps using 26-connectivity labelling (60 Adult-ward clusters, 50 Adolescent-ward clusters) and were fixed before permutation testing. The null distribution was constructed by pooling cluster-level mean betas from both contrast directions and randomly reassigning them to two groups (preserving original group sizes) over 10,000 permutations; *P*-values were computed as (count + 1)/(n_perm + 1). This indicates larger task-general reconfiguration during switches to the typically less-foregrounded Adult. Full results are in Tables S6-S7.

**Figure 3.**
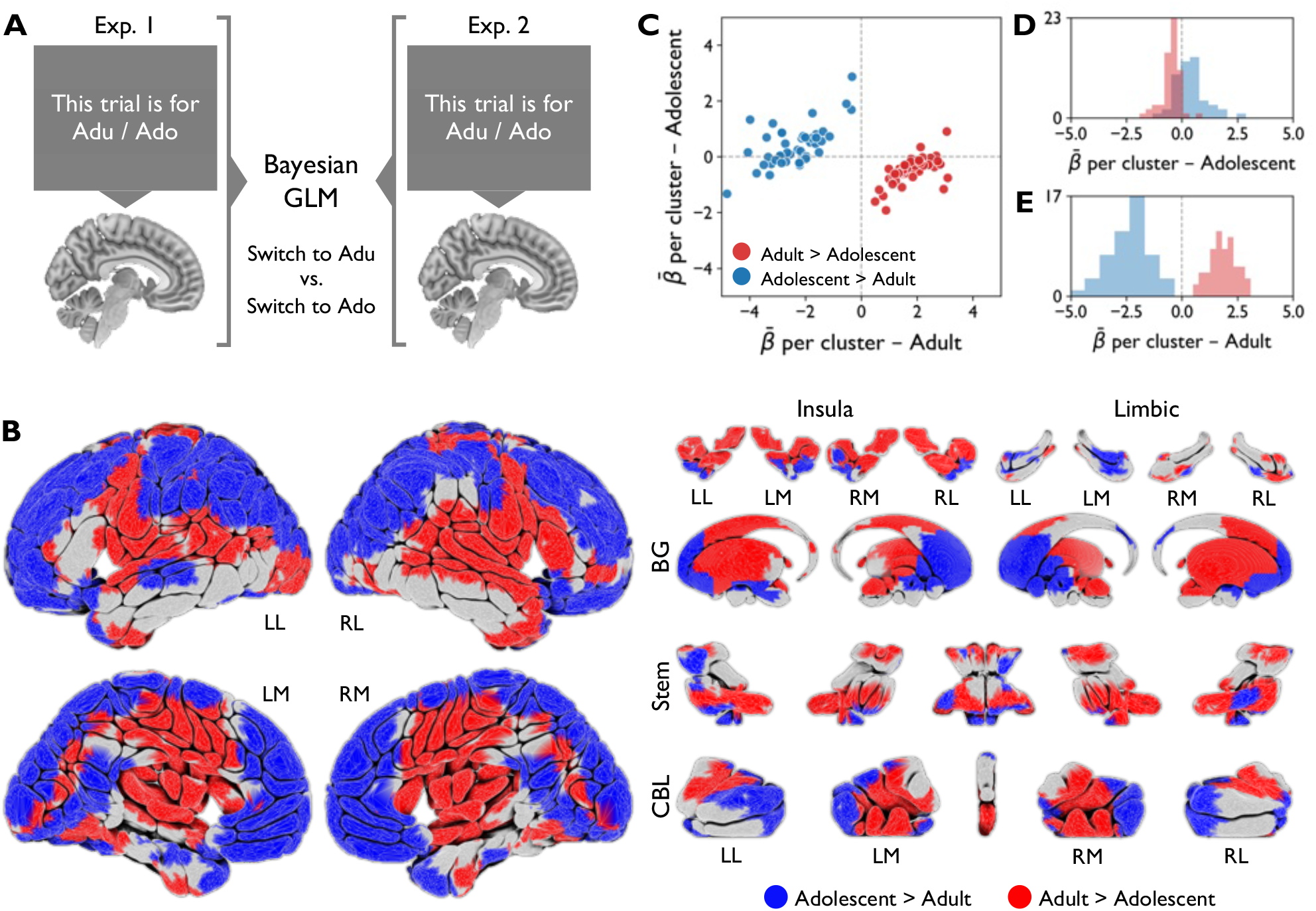
Methods and results of the identity-instruction cue epoch (task-general switching analysis). (A) Schematic of the pooled cue-epoch analysis. Identity-instruction periods (‘Adult’ versus ‘Adolescent’ cues) from the preference/aversion task (Experiment 1) and the trait-judgment task (Experiment 2) were concatenated and modelled in a single Bayesian first-level GLM, with cue events coded by direction of switch (switch to Adult versus switch to Adolescent) independently of task. (B) Surface rendering of Bayesian GLM contrasts for cue-locked activity. Posterior probability maps (PPMs) show regions where BOLD responses during switches to Adult exceeded switches to Adolescent and vice versa, thresholded at the single-case Bayesian level (PPM log-odds >= 10, extent threshold: 20 voxels). (C) Cluster-wise parameter estimates for the two switch directions. Each point represents one significant cluster from (B), plotted by its median beta estimate for switches to Adult (x-axis) and switches to Adolescent (y-axis); accompanying marginal histograms depict the distribution of cluster-level effects for each direction, summarizing the asymmetry in cue-locked modulation between Adult-ward and Adolescent-ward switching. Adu: Adult, Ado: Adolescent, LL: left-lateral view, RL: right-lateral view, LM: left-medial view, RM: right-medial view, BG: basal ganglia, Stem: brainstem, CBL: cerebellum.

### Cross-task identity-invariant judgment analysis

In Experiment 1, separating the perceptual judgment period from picture viewing yielded judgment greater than viewing effects for both identities, and a conjunction identified voxels reliably expressed in both states (Fig. 4). In Experiment 2, a rater-invariant target representation map was derived by retaining target-related structure while excluding rater-dominated structure. Intersecting these two identity-invariant maps isolated a constrained overlap localized primarily to posterior visual and dorsal parietal cortex (Fig. 4), including calcarine/cuneus regions and a superior parietal locus (Table S8). This cross-task overlap provides a conservative estimate of shared task-related neural components preserved across identity-states alongside the identity-selective valuation, person-knowledge and switching signatures.

**Figure 4.**
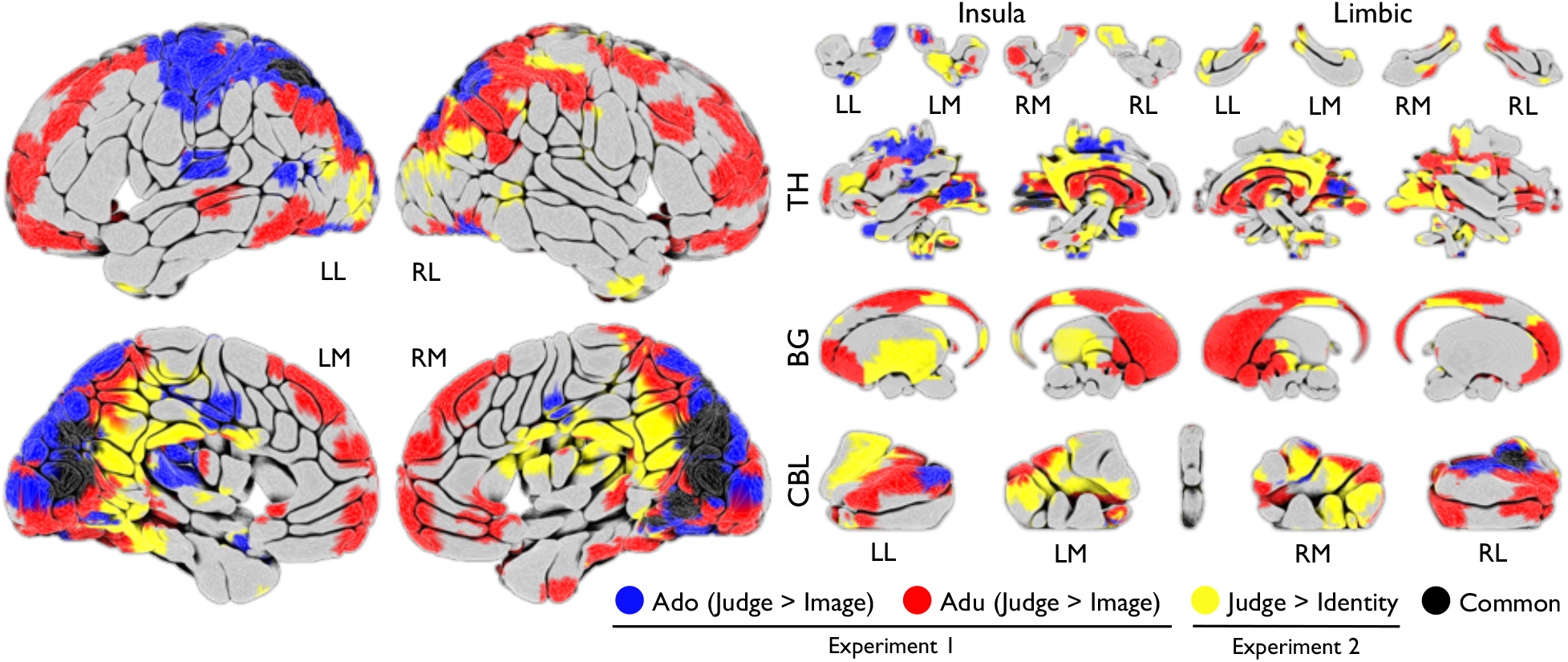
Identity-invariant judgment-related components across experiments (cross-task identity-invariant judgment analysis). Surface renderings show Experiment 1 Bayesian GLM contrasts for the perceptual judgment period relative to picture viewing (Judge > Image) separately for Adolescent (blue) and Adult (red), and the Experiment 2 rater-invariant ‘Judge > Identity’ map (yellow) derived from multi-regression searchlight RSA by retaining voxels with significant Target identity and/or Self-versus-Non-Self effects while excluding voxels with significant Rater identity and/or Raters x Targets effects. Black indicates the overlap between the Experiment 2 Judge > Identity map and voxels significant for Judge > Image in both identity-states in Experiment 1. Bayesian PPMs were thresholded at effect size threshold for PPM: 0.10%, log-odds threshold for PPM: 10, extent threshold: 20 voxels; RSA maps were thresholded at permutation-based voxel-wise *p* < 0.001 with cluster extent >= 20 voxels. Adu: Adult, Ado: Adolescent, LL: left-lateral view, RL: right-lateral view, LM: left-medial view, RM: right-medial view, TH: thalamus, BG: basal ganglia, CBL: cerebellum.

## Discussion

This proof-of-concept study asked whether co-conscious DID, a presentation in which state differences should be hardest to detect, would still show reproducible neural signatures under non-traumatic conditions. The same insect stimuli engaged different valuation-related ensembles and task-evoked coupling in the two identity-states; within canonical self-referential regions, Adult-as-self and Adolescent-as-self occupied distinct self-like positions in person-knowledge space; and switching showed a direction-dependent asymmetry, with larger reconfiguration when switching to the typically less-foregrounded Adult. Cross-task overlap further identified a limited shared task-related substrate in posterior visual and dorsal parietal cortex. The overall pattern is therefore one of selective reconfiguration within a shared neural system, rather than a global change in task engagement.

A practical strength of the ID-SCAN approach is that it does not require traumatic autobiographical recall. Because symptom-provocation paradigms can confound identity-state effects with processes specific to traumatic remembering itself,^15–17,27^ a non-traumatic approach offers a safer and more standardizable route to mechanistic study of DID. That matters clinically as well as experimentally: it suggests that dissociative state expression and switching may be studied without asking participants to relive highly distressing material, thereby expanding population access beyond the subset of patients who can safely tolerate trauma provocation.

The Experiment 1 valuation profiles were not simply mirror images of one another, but engaged qualitatively different neural architectures. The Adolescent-selective insect responses centred on caudate/putamen and mediodorsal thalamus, a circuit associated with habitual or stimulus-driven valuation^39,40^, with gPPI coupling extending to insula and parahippocampal cortex, suggesting viscero-affective and memory-mediated appraisal. By contrast, the Adult-selective profile spanning amygdala, orbitofrontal/medial prefrontal cortex and insula, with broad coupling to lateral prefrontal and temporo-occipital regions, is more consistent with aversion-tagged, top-down evaluative processing.^40,41^ This dissociation maps onto the distinction between reflexive-affective and deliberative-appraisal valuation architectures identified in neuroeconomics research,^40^ and suggests that the two identity-states instantiate not merely different preference ratings but different modes of evaluating the same stimuli. Although reverse inference is limited in a single case,^42^ these coordinated regional and connectivity differences are difficult to explain as a uniform response bias or trial-by-trial acting. The topographic profile of the Adult-selective valuation network, particularly bilateral amygdala and medial prefrontal cortex involvement, shows convergence with the emotion-regulation patterns reported in the apparently normal part (ANP) of DID patients in the PET study by Reinders *et al*.,^7^ while the Adolescent-selective striato-thalamic profile partially maps onto the more habitual or reactive circuitry described for emotional parts (EP) in prior work.^8,43^ This convergence across different paradigms (traumatic recall versus benign valuation) and different imaging modalities (PET versus fMRI) strengthens the case that identity-state-dependent neural organization is not an artefact of trauma-provocation procedures.

Experiment 2 strengthened this point in a more abstract domain. The key result was not just that ratings differed, but that each identity occupied a distinct self-like position within canonical self-referential network. Within the cortical midline structures implicated in self-referential processing,^45,46^ the rater identity model produced the broadest footprint, consistent with a widespread reconfiguration of person-knowledge representations when the foreground identity changed, while target-related structure was most prominent in precuneus, a region linked to episodic self-knowledge and autobiographical memory.^46^ From a predictive processing perspective, the fact that both Adult-as-self and Adolescent-as-self positions were self-like (i.e. closer to each other than to non-self judgments) could be interpreted as two distinct prior self-models instantiated within overlapping cortical populations, with identity-cued switching selecting among attractor states rather than rebuilding self-representation de novo.^47,48^ The shift in representation of the same intimate other depending on who was judging further suggests that person knowledge is reorganized by the rater’s identity, not merely relabelled.

The pooled cue analysis extended the findings from state representation to state transition. Larger cue-locked reconfiguration when switching to Adult suggests direction-dependent switching costs in this case. This asymmetry is consistent with the broader task-switching literature, where switch costs are systematically larger for the less-practised or less-dominant task set,^49,50^ and with clinical descriptions of dominance asymmetry in DID.^6,12^ The history that Adolescent usually managed daily interactions implies that Adult represents the less-automatized state. The regions showing greater modulation for Adult-ward switches (medial frontal/cingulate, insula, brainstem) overlap with the conflict-monitoring and task-set reconfiguration networks identified in cognitive task-switching neuroimaging.^49^ Earlier fMRI studies of identity switching in DID^13,14^ reported frontal and temporal involvement but lacked the task-general pooling and quantitative asymmetry indices employed here.

The cross-task overlap analysis complements these identity-selective findings by showing that some task-related processing remained shared across identities. The localization of this overlap to posterior visual and dorsal parietal cortex is consistent with domain-general perceptual encoding and attention/working-memory demands shared by any two visual cognitive tasks,^24–26^ while the absence of prefrontal and self-referential regions from the invariant map is notable: it suggests that higher-order integrative processing is more identity-state-dependent than low-level perceptual encoding, a pattern of selective reconfiguration within a shared neural system, rather than a global change in task engagement. This internal baseline constrains interpretation by showing that marked identity-selective effects coexisted with preserved perceptual and attentional resources for carrying out the tasks.

These convergent findings speak directly to the concern that DID reflects deliberate enactment rather than psychobiological organization.^1–3,9,10^ Prior studies using instructed simulators demonstrated that healthy individuals could not reproduce the identity-state-dependent PET and fMRI activation patterns of genuine DID patients when asked to simulate multiple identities in response to traumatic scripts.^9,10^ The present study provides a complementary, structurally distinct test. Although it did not include a formal simulator group, the ID-SCAN design offers several features that are difficult to reconcile with voluntary strategic performance: (i) the use of identical stimuli across identity-cued conditions, so that any neural difference cannot arise from stimulus properties; (ii) the trial-by-trial identity-check screen providing a real-time compliance measure; (iii) the representational geometry result, which requires not only different response patterns but a specific self-like organizational structure that would be difficult to strategically produce; and (iv) the convergence across analytically independent methods (activation, connectivity and representational similarity), which multiplies the constraints that any simulation account must simultaneously satisfy. Under a pure acting account, especially in a co-conscious presentation, one would expect weak or inconsistent task effects; instead, we observed coherent, convergent identity-selective signatures alongside a limited shared task-related baseline. We acknowledge, however, that co-consciousness also permits voluntary access to both perspectives, meaning that the present design cannot definitively rule out sophisticated strategic performance. Future studies should include instructed-simulator and clinical comparison groups to formally test specificity.

A key design feature of ID-SCAN is that it separates fixed structural components (trial architecture, identity-cue epoch design, Bayesian single-case statistics and cross-task overlap analysis) from participant-calibrated components that are individually tailored during a pre-scan clinical interview: stimulus selection for the valuation task (here, insects) and the choice of intimate other for the trait-judgment task. This modular structure means the protocol can be applied to other DID cases by substituting stimuli identified through the same semi-structured interview process, without modifying the analytical framework. If replicated, the most tractable candidate metrics for longitudinal treatment-tracking include the switching-asymmetry ratio (Adult-ward versus Adolescent-ward cue-locked beta separation), the Euclidean distance between self-positions in the RSA geometry, and the spatial extent of the cross-task identity-invariant overlap. Treatment-related integration would be predicted to reduce the switching-asymmetry ratio (as the less-foregrounded identity becomes more accessible), decrease the RSA distance between self-positions (as identity-state representations converge), and potentially expand the identity-invariant overlap (as more processing becomes shared). These predictions are testable in longitudinal single-case and cohort designs.

This report concerns a single co-conscious DID case, and the absence of trauma-exposed controls, clinical comparison groups and instructed simulators limits claims about diagnostic specificity. Although the ID-SCAN design provides structural protections against simulation (identical stimuli, identity-check screen, multi-method convergence), a formal simulator comparison analogous to those conducted by Reinders *et al*.^9^ and Vissia *et al*.^10^ remains necessary to establish that the observed neural patterns cannot be produced by healthy individuals instructed to adopt alternating cognitive perspectives. The inclusion of such a comparison group was not feasible in the present proof-of-concept study but is planned for subsequent work.

The preference task used idiosyncratic but clinically meaningful stimuli, and the trait-judgment geometry may partly depend on the selected intimate other and cultural context. As a single-case study, all findings are potentially specific to this individual’s history, personality and clinical configuration; generalizability to other DID presentations, including amnestic or polyfragmented cases, remains to be tested. The study also provides no information about the theory of structural dissociation of the personality (TSDP)^6,44^ in the formal sense; that is, whether the Adult and Adolescent identity-states correspond to apparently normal and emotional parts as defined in that framework cannot be assessed because the TSDP’s predictions about differential physiological reactivity and somatic experience were not assessed.

Despite these limitations, single-case methodology has a longstanding and productive role in cognitive neuroscience,^37,38^ and the present design provides internal replication across two tasks and a pooled cue analysis that together constitute multiple within-person tests of the same underlying hypothesis. The data show that a non-traumatic experimental protocol can detect coherent neural signatures of identity-state organization and switching under co-conscious conditions, motivating the cohort and longitudinal studies needed to establish specificity and clinical utility.

## Data availability

The data that support the findings of this study are available from the corresponding author, S.K., upon reasonable request. The data are not publicly available because they contain information that could compromise participant privacy. Analysis code for all preprocessing, modelling, statistical inference and visualization steps reported in this study is deposited at [OSF/Zenodo URL to be inserted upon acceptance] and includes a README documenting the execution order and software dependencies. De-identified derived outputs will be shared in line with privacy constraints.

## Acknowledgements

We thank the participant for her time and willingness to engage in the identity-state experiments, and we appreciate the support of the IRCN Human fMRI Core, the University of Tokyo Institutes for Advanced Studies. During analysis and manuscript preparation, the authors used ChatGPT (OpenAI, October-December 2025) and Claude Code (Anthropic, March-April 2026) as writing and coding assistants. All outputs were reviewed, verified and revised by the authors, who take full responsibility for the final content.

## Supplementary material

Supplementary material is available at *Brain Communications* online.

## Author contributions

Conception and design of the study: S.K. Task design (ID-SCAN protocol and experimental paradigms): S.K., A.H., and K.O. MRI protocol development and optimization: S.K. and R.N. Participant recruitment and clinical assessment: K.O. MRI data acquisition: S.K., J.Y., H.F. and A.I. Behavioural data collection: S.K. Data analysis and figure preparation: S.K. Data interpretation: S.K., N.A., S.N., K.O. and T.M. Drafting of the manuscript: S.K. Critical revision of the manuscript for important intellectual content: N.A., S.N., K.O., A.I. and T.M. Supervision: T.M. Funding acquisition and project administration: S.K. Final approval of the version to be published: all authors.

