## Supplemental Tables for "Non-traumatic fMRI of identity-state expression and switching in co-conscious DID"

Table S1. Bayesian GLM main effect of stimulus (insects > scrambled) in Experiment 1.

| Region | [x, y, z] | Cl. size | Log odds | Region | [x, y, z] | Cl. Size | Log odds |
|---|---|---|---|---|---|---|---|
| ACC_sup | [2, 16, 20] | 29 | 23.72 | Olfactory | [0, -42, 74] | 22 | 17.46 |
| Angular | [-44, -66, 48] | 21 | 23.88 | Paracentral_Lobule | [-44, -28, 36] | 45 | 36.04 |
| Angular | [44, -52, 28] | 38 | 24.21 | Parietal_Inf | [12, -38, 82] | 22 | 19.51 |
| Caudate | [18, 10, 20] | 25 | 17.64 | Postcentral | [-36, 0, 28] | 109 | 36.04 |
| Caudate | [18, -6, 22] | 28 | 16.05 | Precentral | [-32, -2, 44] | 1656 | 36.04 |
| Cerebellum_10 | [-18, -28, -42] | 33 | 35.13 | Precentral | [-50, 8, 40] | 47 | 26.10 |
| Cerebellum_3 | [22, -26, -30] | 37 | 31.11 | Precentral | [48, 4, 32] | 25 | 22.23 |
| Cerebellum_6 | [22, -70, -26] | 22 | 17.74 | Precentral | [46, -12, 42] | 21 | 16.77 |
| Cerebellum_6 | [34, -66, -24] | 33 | 18.56 | Precentral | [-2, -56, 18] | 1836 | 36.04 |
| Cerebellum_8 | [-36, -44, -56] | 86 | 36.04 | Precuneus | [2, -46, 64] | 34 | 27.84 |
| Cerebellum_9 | [6, -60, -50] | 25 | 20.92 | Precuneus | [-2, 0, 64] | 63 | 23.46 |
| Cerebellum_Crus1 | [-58, -48, -36] | 22 | 32.98 | Supp_Motor_Area | [-62, -46, 30] | 21 | 18.71 |
| Cerebellum_Crus1 | [-48, -58, -36] | 22 | 13.54 | SupraMarginal | [62, -32, 44] | 31 | 36.04 |
| Cerebellum_Crus1 | [-44, -74, -28] | 8607 | 36.04 | SupraMarginal | [-54, 12, -6] | 275 | 35.35 |
| Cerebellum_Crus1 | [28, -88, -28] | 23 | 25.82 | Temporal_Pole_Sup | [38, 14, -22] | 117 | 36.04 |
| Frontal_Inf_Oper | [-44, 14, 6] | 57 | 18.03 | Temporal_Pole_Sup | [64, 2, 2] | 473 | 36.04 |
| Frontal_Inf_Oper | [44, 4, 20] | 406 | 36.04 | Temporal_Sup | [62, -50, 18] | 21 | 21.19 |
| Frontal_Inf_Orb_2 | [48, 44, -10] | 86 | 21.59 | Temporal_Sup | [8, -28, 8] | 822 | 36.04 |
| Frontal_Inf_Tri | [-34, 24, 16] | 132 | 32.23 | Thal_PuM | [4, -36, -18] | 37 | 30.48 |
| Frontal_Med_Orb | [10, 42, -10] | 21 | 19.06 | Vermis_1_s2 | [0, -64, 0] | 27 | 20.94 |
| Frontal_Mid_2 | [36, 34, 10] | 138 | 36.04 | Vermis_4_5 | [-4, -34, -60] | 131 | 36.04 |
| Frontal_Mid_2 | [32, 52, 34] | 59 | 36.04 | ANGULAR_WM | [-24, -50, 38] | 133 | 36.04 |
| Frontal_Sup_2 | [-22, 60, -2] | 92 | 25.90 | CEREBELLUM | [-12, -28, -22] | 30 | 30.99 |
| Frontal_Sup_2 | [-16, 36, 42] | 45 | 26.75 | CT | [-20, 18, -12] | 74 | 21.63 |
| Frontal_Sup_2 | [22, 46, 30] | 32 | 20.42 | IFOF | [-16, -26, 24] | 22 | 16.52 |
| Frontal_Sup_2 | [20, 24, 46] | 23 | 20.66 | MCP | [2, -42, -28] | 62 | 36.04 |
| Frontal_Sup_2 | [24, -8, 52] | 22 | 22.19 | MEDULLA | [12, -32, -52] | 701 | 36.04 |
| Frontal_Sup_Medial | [10, 62, 4] | 910 | 36.04 | MEDULLA | [0, -42, 74] | 22 | 17.46 |
| Heschl | [42, -20, 12] | 27 | 23.81 | SPWM | [0, -56, 76] | 22 | 16.63 |
| Insula | [-40, 10, -14] | 48 | 24.44 | | [36, -44, 34] | 58 | 36.04 |
| Insula | [44, 10, -14] | 75 | 36.04 | | [22, -8, -46] | 65 | 33.18 |
| Lingual | [10, -34, -2] | 358 | 36.04 | | [-18, -28, -32] | 44 | 28.84 |
| Occipital_Mid | [-30, -84, 18] | 549 | 36.04 | | | | |

Lowercase region labels denote AAL; uppercase labels denote JHU.

CT: CORTICOSPINAL_TRACT; INFERIOR_FRONT_OCCIPITAL_FASCICULUS: IFOF; MCP: MIDDLE_CEREBELLAR_PEDUNCLE; SPWM: SUPERIOR_PARIETAL_WM.

Table S2. Bayesian GLM main effect of identity-state (Adolescent > Adult) in Experiment 1.

| Region | [x, y, z] | Cl. size | Log odds |
|---|---|---|---|
| Frontal_Sup_2 | [-14, 56, 24] | 241 | 35.13 |
| Frontal_Sup_2 | [28, 60, 12] | 71 | 34.43 |
| Frontal_Sup_2 | [26, 44, 44] | 89 | 19.64 |
| Frontal_Sup_Medial | [10, 56, 20] | 61 | 20.64 |
| Paracentral_Lobule | [-8, -36, 68] | 33 | 20.06 |
| Paracentral_Lobule | [4, -24, 74] | 24 | 20.13 |
| Postcentral | [52, -30, 60] | 44 | 30.24 |
| Postcentral | [12, -42, 72] | 42 | 20.24 |
| Supp_Motor_Area | [-8, -6, 74] | 20 | 18.73 |
| Temporal_Pole_Sup | [-36, 8, -26] | 95 | 36.04 |
| MIDBRAIN | [2, -10, -8] | 22 | 19.80 |
| FORNIX | [2, 6, 8] | 47 | 36.04 |
| | [26, -8, -48] | 22 | 36.04 |
| | [2, -42, -30] | 23 | 24.74 |

Lowercase region labels denote AAL; uppercase labels denote JHU.

Table S3. Bayesian GLM main effect of identity-state (Adult > Adolescent) in Experiment 1.

| Region | [x, y, z] | Cl. size | Log odds | Region | [x, y, z] | Cl. size | Log odds |
|---|---|---|---|---|---|---|---|
| Angular | [-50, -62, 44] | 24 | 17.93 | Occipital_Sup | [26, -98, 12] | 23 | 17.29 |
| Angular | [48, -66, 44] | 34 | 24.59 | OFCmed | [-14, 26, -20] | 55 | 24.08 |
| Caudate | [14, -2, 16] | 25 | 18.98 | Paracentral_Lobule | [-14, -18, 80] | 29 | 22.74 |
| Cerebellum_3 | [16, -30, -18] | 29 | 26.39 | ParaHippocampal | [-14, -28, -22] | 32 | 18.90 |
| Cerebellum_4_5 | [30, -32, -36] | 45 | 30.94 | Postcentral | [-48, -12, 30] | 897 | 36.04 |
| Cerebellum_6 | [32, -50, -32] | 48 | 24.82 | Precentral | [-44, 12, 30] | 27 | 23.05 |
| Cerebellum_8 | [-38, -44, -54] | 84 | 36.04 | Precuneus | [0, -54, 48] | 25 | 21.55 |
| Cerebellum_8 | [-34, -46, -44] | 71 | 32.41 | Putamen | [-22, 18, -10] | 197 | 28.56 |
| Cerebellum_8 | [18, -74, -52] | 22 | 23.78 | Rectus | [10, 24, -14] | 29 | 16.15 |
| Cerebellum_9 | [10, -62, -52] | 21 | 25.77 | SupraMarginal | [66, -28, 38] | 22 | 26.91 |
| Cerebellum_Crus1 | [-28, -86, -24] | 2171 | 36.04 | Temporal_Mid | [-64, -10, -6] | 1050 | 36.04 |
| Cingulate_Mid | [0, -4, 44] | 292 | 36.04 | Temporal_Mid | [-60, -60, 4] | 211 | 36.04 |
| Cingulate_Mid | [2, 20, 32] | 23 | 14.55 | Temporal_Mid | [68, -10, -12] | 5779 | 36.04 |
| Cingulate_Post | [2, -42, 20] | 259 | 36.04 | Temporal_Pole_Mid | [-34, 14, -32] | 25 | 21.94 |
| Cuneus | [-8, -102, 16] | 28 | 17.07 | Temporal_Pole_Sup | [-36, 12, -22] | 36 | 25.05 |
| Frontal_Inf_Orb_2 | [-42, 18, -12] | 23 | 25.94 | Temporal_Pole_Sup | [40, 12, -20] | 68 | 25.33 |
| Frontal_Inf_Orb_2 | [48, 46, -6] | 62 | 19.07 | Thal_VA | [-4, -2, 4] | 34 | 20.29 |
| Frontal_Mid_2 | [30, 60, 26] | 31 | 24.44 | Thal_VA | [-6, -4, 12] | 46 | 23.44 |
| Frontal_Sup_2 | [-28, 60, -2] | 50 | 28.51 | Vermis | [2, -42, -34] | 103 | 25.18 |
| Frontal_Sup_2 | [28, 60, 6] | 795 | 36.04 | Vermis | [2, -44, -6] | 37 | 33.24 |
| Heschl | [40, -26, 16] | 22 | 25.46 | CEREBELLUM | [0, 6, -18] | 54 | 25.82 |
| Insula | [-38, -12, 16] | 22 | 24.89 | CEREBELLUM | [-10, -74, -54] | 69 | 29.09 |
| Lingual | [14, -32, -8] | 38 | 18.71 | MCP | [-18, -28, -40] | 64 | 24.72 |
| Lingual | [6, -36, 2] | 148 | 36.04 | MCP | [-16, -26, -30] | 57 | 34.43 |
| Occipital_Inf | [32, -90, -6] | 25 | 22.28 | | [18, -16, -30] | 27 | 19.86 |
| Occipital_Sup | [-16, -84, 48] | 51 | 36.04 | | [10, -38, -62] | 461 | 36.04 |

Lowercase region labels denote AAL; uppercase labels denote JHU.

MCP: MIDDLE_CEREBELLAR_PEDUNCLE.

Table S4. Bayesian GLM identity-selective spotlight contrast (Adolescent-insect > others) and corresponding gPPI connectivity in Experiment 1.

| Bayesian GLM | | | | gPPI | | | |
|---|---|---|---|---|---|---|---|
| Region | [x, y, z] | Cl. size | Log odds | Region | [x, y, z] | Cl. size | Peak Stat |
| Caudate | [14, 22, 4] | 33 | 9.09 | Insula | [-46, 2, 0] | 33 | 5.85 |
| | | | | ParaHippocampal | [20, -22, -16] | 26 | 5.91 |
| Putamen | [-18, 12, 8] | 20 | 7.95 | Fusiform | [26, -82, -16] | 21 | 4.33 |
| | | | | Precentral | [-36, -20, 66] | 25 | 4.51 |
| | | | | Precentral | [34, -16, 70] | 30 | 5.30 |
| | | | | Precuneus | [4, -72, 44] | 22 | 4.64 |
| Thal_MDm | [-4, -22, 8] | 97 | 18.05 | SupraMarginal | [56, -42, 42] | 74 | 5.79 |
| | | | | Frontal_Sup_2 | [28, 60, 6] | 27 | 5.05 |
| | | | | Precentral | [30, -20, 70] | 24 | 4.42 |

Region labels denote AAL.

Table S5. Bayesian GLM identity-selective spotlight contrast (Adult-insect > others) and corresponding gPPI connectivity in Experiment 1.

| Bayesian GLM | | | | gPPI | | | |
|---|---|---|---|---|---|---|---|
| Region | [x, y, z] | Cl. size | Log odds | Region | [x, y, z] | Cl. size | Peak Stat |
| Amygdala_R | [30, 4, -18] | 46 | 23.76 | Angular | [-40, -62, 28] | 46 | 5.72 |
| | | | | Frontal_Sup_Medial | [-12, 60, 10] | 57 | 5.34 |
| | | | | Temporal_Inf | [52, -62, -8] | 28 | 5.00 |
| | | | | Temporal_Mid | [68, -22, -16] | 32 | 6.12 |
| | | | | Temporal_Mid | [68, -46, 2] | 35 | 5.44 |
| Frontal_Inf_Orb_2_L | [-44, 38, -12] | 101 | 12.96 | Amygdala | [24, 0, -14] | 39 | 6.10 |
| | | | | Angular | [42, -72, 40] | 22 | 5.87 |
| | | | | Calcarine | [20, -92, -4] | 31 | 5.39 |
| | | | | Calcarine | [4, -74, 16] | 24 | 5.37 |
| | | | | Caudate | [-6, 0, 14] | 33 | 6.05 |
| | | | | Cerebellum_3 | [20, -28, -28] | 22 | 4.93 |
| | | | | Cerebellum_Crus1 | [50, -66, -22] | 26 | 6.09 |
| | | | | Cerebellum_Crus1 | [30, -86, -30] | 29 | 5.48 |
| | | | | Cerebellum_Crus1 | [8, -90, -22] | 27 | 5.24 |
| | | | | Cingulate_Mid | [0, -6, 46] | 34 | 5.33 |
| | | | | Cuneus | [-6, -88, 36] | 48 | 5.87 |
| | | | | Frontal_Sup_2 | [24, 68, 18] | 44 | 6.76 |
| | | | | Frontal_Sup_Medial | [-6, 66, 16] | 79 | 6.25 |
| | | | | Fusiform | [38, -76, -18] | 43 | 6.71 |
| | | | | Hippocampus | [-14, -8, -22] | 32 | 6.24 |
| | | | | Insula | [46, 2, -8] | 44 | 6.32 |
| | | | | Lingual | [14, -68, -10] | 21 | 5.30 |
| | | | | Lingual | [20, -94, -10] | 29 | 4.91 |
| | | | | Occipital_Inf | [34, -92, -14] | 126 | 6.60 |
| | | | | Occipital_Mid | [34, -82, 32] | 86 | 7.19 |
| | | | | Occipital_Mid | [30, -96, 8] | 25 | 5.22 |
| | | | | Occipital_Sup | [-12, -102, 10] | 43 | 5.73 |
| | | | | Occipital_Sup | [24, -84, 42] | 22 | 5.36 |
| | | | | Olfactory | [8, 14, -16] | 40 | 5.81 |
| | | | | Paracentral_Lobule | [-6, -28, 76] | 41 | 5.49 |
| | | | | ParaHippocampal | [22, 4, -20] | 26 | 6.52 |
| | | | | Parietal_Sup | [-18, -70, 56] | 90 | 6.58 |
| | | | | Parietal_Sup | [18, -78, 52] | 71 | 6.56 |

| | | | | | | | |
|---|---|---|---|---|---|---|---|
| | | | | Postcentral | [16, -44, 78] | 88 | 6.52 |
| | | | | Precentral | [34, -26, 74] | 23 | 5.85 |
| | | | | Precuneus | [0, -68, 60] | 70 | 6.71 |
| | | | | Precuneus | [4, -46, 62] | 21 | 6.50 |
| | | | | Precuneus | [4, -74, 42] | 29 | 5.23 |
| | | | | SupraMarginal | [54, -24, 28] | 23 | 6.04 |
| | | | | Temporal_Inf | [50, -58, -26] | 33 | 5.85 |
| | | | | Temporal_Mid | [-62, -34, -16] | 22 | 5.03 |
| | | | | Temporal_Mid | [50, -66, 14] | 22 | 6.26 |
| | | | | Temporal_Mid | [52, -62, 0] | 28 | 5.07 |
| | | | | Vermis_3 | [6, -40, -4] | 44 | 5.80 |
| | | | | CEREBELLUM | [-12, -46, -62] | 20 | 5.23 |
| | | | | ML | [-6, -40, -38] | 29 | 6.23 |
| | | | | SCP | [-10, -28, -18] | 20 | 5.19 |
| | | | | | [16, -28, -46] | 21 | 6.71 |
| | | | | | [2, -26, 12] | 23 | 4.65 |
| Frontal_Inf_Orb_2_R | [48, 50, -4] | 940 | 29.11 | Frontal_Sup_2 | [-18, 64, 4] | 46 | 6.67 |
| | | | | Frontal_Sup_2 | [28, 66, 8] | 22 | 4.79 |
| | | | | Frontal_Sup_Medial | [-8, 62, 2] | 20 | 5.44 |
| | | | | | [2, 6, -18] | 24 | 4.46 |
| Frontal_Med_Orb_R | [16, 56, -6] | 420 | 15.18 | Frontal_Med_Orb | [-8, 64, -10] | 25 | 4.68 |
| | | | | Frontal_Sup_2 | [18, 64, 8] | 94 | 6.92 |
| Insula_L | [-46, -10, 4] | 1024 | 28.27 | | [2, 6, -18] | 24 | 5.97 |
| Putamen_R | [24, 20, -2] | 86 | 19.12 | Cerebellum_6 | [6, -86, -16] | 43 | 5.63 |
| | | | | Cerebellum_Crus1 | [-30, -84, -24] | 66 | 5.33 |
| | | | | Cerebellum_Crus2 | [-54, -48, -48] | 22 | 5.47 |
| | | | | Cuneus | [2, -86, 34] | 20 | 4.80 |
| | | | | Frontal_Med_Orb | [-6, 62, -4] | 25 | 5.87 |
| | | | | Precuneus | [-2, -40, 80] | 22 | 5.08 |
| Thal_MDm_L | [-4, -18, 0] | 35 | 19.93 | Frontal_Sup_2 | [-22, 60, 0] | 122 | 6.76 |
| | | | | Frontal_Sup_2 | [18, 58, 4] | 21 | 5.75 |
| Thal_MDm_R | [4, -18, 0] | 37 | 36.04 | Frontal_Med_Orb | [2, 58, -12] | 20 | 7.23 |
| | | | | Frontal_Sup_Medial | [12, 72, 10] | 437 | 8.01 |
| Thal_VA_R | [8, 0, 4] | 480 | 36.04 | Frontal_Sup_Medial | [-2, 52, 44] | 43 | 6.16 |
| | | | | Heschl | [-66, -8, 8] | 22 | 6.55 |
| | | | | | [2, 12, -22] | 28 | 7.22 |

Lowercase region labels denote AAL; uppercase labels denote JHU.

SCP: SUPERIOR_CEREBELLAR_PEDUNCLE; ML: MEDIAL_LEMNISCUS.

Table S6. Bayesian GLM identity-switching contrast (switches to Adolescent > switches to Adult) in task-general switching analysis.

| Region | [x, y, z] | Cl. size | Log odds | Region | [x, y, z] | Cl. size | Log odds |
|---|---|---|---|---|---|---|---|
| Angular | [56, -58, 32] | 26 | 27.09 | OFCant | [20, 44, -14] | 52 | 27.92 |
| Angular | [48, -64, 48] | 50 | 36.04 | OFCpost | [-38, 22, -16] | 32 | 28.29 |
| Calcarine | [-4, -102, 2] | 290 | 36.04 | OFCpost | [24, 24, -20] | 296 | 36.04 |
| Calcarine | [22, -102, 0] | 477 | 36.04 | Paracentral_Lobule | [4, -28, 78] | 23 | 27.09 |
| Caudate | [4, 10, -2] | 502 | 36.04 | Parietal_Sup | [30, -46, 46] | 26 | 24.59 |
| Caudate | [18, 26, 4] | 26 | 27.66 | Parietal_Sup | [54, -34, 58] | 35 | 23.30 |
| Cerebellum_4_5 | [28, -28, -32] | 48 | 36.04 | Postcentral | [-64, -14, 34] | 29 | 27.63 |
| Cerebellum_Crus2 | [-20, -84, -44] | 81 | 36.04 | Postcentral | [-52, -28, 56] | 94 | 36.04 |
| Cerebellum_Crus2 | [24, -86, -42] | 852 | 36.04 | Postcentral | [-42, -16, 52] | 25 | 26.69 |
| Frontal_Mid_2 | [46, 18, 44] | 67 | 33.40 | Postcentral | [-36, -30, 66] | 26 | 31.07 |
| Frontal_Mid_2 | [44, 30, 42] | 20 | 22.86 | Precentral | [52, -12, 42] | 108 | 36.04 |
| Frontal_Mid_2 | [36, 8, 58] | 21 | 20.17 | Precentral | [38, -26, 58] | 25 | 21.33 |
| Frontal_Sup_2 | [30, -14, 60] | 30 | 24.59 | Precentral | [14, -24, 76] | 22 | 22.45 |
| Frontal_Sup_2 | [12, -10, 76] | 22 | 35.35 | Precuneus | [0, -56, 12] | 116 | 35.13 |
| Frontal_Sup_Medial | [10, 50, 6] | 31 | 21.54 | Precuneus | [4, -68, 48] | 29 | 36.04 |
| Frontal_Sup_Medial | [14, 30, 62] | 68 | 30.29 | Rectus | [6, 22, -22] | 581 | 36.04 |
| Fusiform | [24, 6, -46] | 49 | 36.04 | Supp_Motor_Area | [-4, -2, 78] | 152 | 36.04 |
| Hippocampus | [-22, -10, -26] | 21 | 18.51 | SupraMarginal | [-64, -40, 38] | 40 | 29.63 |
| Hippocampus | [-22, -12, -16] | 46 | 25.08 | Temporal_Mid | [-66, -28, -6] | 38 | 31.85 |
| Insula | [-24, 14, -20] | 2789 | 36.04 | Temporal_Pole_Mid | [-34, 20, -40] | 45 | 36.04 |
| Lingual | [16, -76, -12] | 21 | 19.24 | Temporal_Pole_Mid | [-34, 24, -36] | 131 | 36.04 |
| Occipital_Mid | [-38, -86, 30] | 554 | 36.04 | Temporal_Pole_Sup | [36, 26, -32] | 43 | 36.04 |
| Occipital_Mid | [44, -76, 14] | 32 | 22.86 | Temporal_Sup | [54, -50, 22] | 21 | 20.00 |
| Occipital_Mid | [34, -88, 26] | 717 | 36.04 | MEDULLA | [-2, -36, -64] | 456 | 36.04 |
| Occipital_Sup | [-10, -84, 48] | 41 | 26.32 | | [20, -36, 22] | 42 | 26.33 |

Lowercase region labels denote AAL; uppercase labels denote JHU.

Table S7. Bayesian GLM identity-switching contrast (switches to Adult > switches to Adolescent) in task-general switching analysis.

| Region | [x, y, z] | Cl. size | Log odds | Region | [x, y, z] | Cl. size | Log odds |
|---|---|---|---|---|---|---|---|
| ACC_pre | [8, 38, 2] | 22 | 26.40 | Occipital_Inf | [40, -68, -16] | 26 | 20.92 |
| Calcarine | [-12, -104, -4] | 1120 | 36.04 | Occipital_Inf | [32, -82, -2] | 414 | 34.95 |
| Calcarine | [-2, -76, 12] | 22 | 25.47 | Occipital_Mid | [-42, -70, 16] | 27 | 22.33 |
| Caudate | [-16, 18, 4] | 29 | 19.71 | Occipital_Mid | [26, -92, 4] | 24 | 24.70 |
| Cerebellum_4_5 | [-20, -40, -24] | 32 | 36.04 | Occipital_Mid | [30, -74, 30] | 81 | 32.12 |
| Cerebellum_6 | [-24, -52, -28] | 739 | 36.04 | Occipital_Sup | [22, -62, 44] | 28 | 22.98 |
| Cerebellum_6 | [24, -52, -34] | 78 | 27.24 | Postcentral | [-22, -36, 74] | 143 | 25.23 |
| Cerebellum_8 | [-14, -70, -52] | 228 | 36.04 | Postcentral | [38, -44, 62] | 36 | 30.30 |
| Cerebellum_8 | [18, -64, -54] | 217 | 36.04 | Postcentral | [26, -40, 62] | 79 | 35.64 |
| Cerebellum_Crus2 | [-4, -80, -42] | 22 | 19.85 | Precuneus | [-12, -50, 48] | 92 | 29.66 |
| Cingulate_Mid | [-8, 12, 32] | 814 | 36.04 | Precuneus | [10, -44, 54] | 51 | 24.58 |
| Cingulate_Mid | [10, -20, 40] | 29 | 21.33 | Precuneus | [10, -60, 58] | 21 | 29.17 |
| Cuneus | [-10, -90, 22] | 20 | 22.43 | Supp_Motor_Area | [-6, -10, 60] | 190 | 36.04 |
| Cuneus | [-8, -70, 24] | 134 | 36.04 | Temporal_Mid | [-64, -8, -6] | 2851 | 36.04 |
| Cuneus | [16, -68, 32] | 169 | 35.35 | Temporal_Mid | [-60, -56, 6] | 232 | 36.04 |
| Frontal_Inf_Oper | [-40, 14, 12] | 24 | 22.94 | Temporal_Mid | [54, -56, 0] | 96 | 36.04 |
| Frontal_Inf_Oper | [38, 14, 28] | 96 | 35.64 | Temporal_Pole_Mid | [-34, 6, -44] | 156 | 33.37 |
| Frontal_Med_Orb | [16, 54, -2] | 37 | 29.60 | Temporal_Pole_Sup | [-34, 12, -30] | 36 | 36.04 |
| Frontal_Mid_2 | [-30, 40, -8] | 25 | 24.46 | Temporal_Pole_Sup | [46, 22, -32] | 3949 | 36.04 |
| Frontal_Mid_2 | [-42, 46, -6] | 44 | 29.39 | Temporal_Sup | [64, -44, 16] | 97 | 36.04 |
| Frontal_Mid_2 | [-36, 46, 4] | 26 | 28.95 | Thal_MDm | [-6, -18, -2] | 90 | 30.51 |
| Frontal_Mid_2 | [36, 44, -12] | 20 | 21.60 | Thal_VL | [-12, -18, 10] | 24 | 27.52 |
| Frontal_Mid_2 | [44, 54, -2] | 23 | 21.68 | Vermis_9 | [0, -58, -38] | 25 | 18.09 |
| Frontal_Sup_2 | [-28, 62, 22] | 30 | 25.10 | MEDULLA | [-6, -44, -64] | 1194 | 36.04 |
| Frontal_Sup_2 | [16, -10, 70] | 26 | 25.83 | IFWM | [34, 40, -4] | 20 | 23.08 |
| Fusiform | [-32, -40, -14] | 20 | 20.23 | PTR | [-22, -58, 18] | 26 | 28.05 |
| Fusiform | [32, -48, -6] | 32 | 32.05 | CINGULUM_WM | [-14, -30, 36] | 51 | 30.17 |
| Insula | [-32, 12, -18] | 39 | 31.05 | POSTCENTRAL_WM | [22, -30, 52] | 266 | 36.04 |
| Insula | [-30, 12, 4] | 168 | 36.04 | POSTCENTRAL_WM | [-20, -32, 52] | 198 | 36.04 |
| Lingual | [-8, -72, -6] | 24 | 21.55 | | [34, -6, -50] | 422 | 36.04 |

Lowercase region labels denote AAL; uppercase labels denote JHU.

IFWM: INFERIOR_FRONTAL_WM, PTR: POSTERIOR_THALAMIC_RADIATION.

Table S8. Overlap of identity-independent judgment-related regions between Experiment 1 and Experiment 2.

| Region | [x, y, z] | Cl. size |
|---|---|---|
| Calcarine | [4, -80, 6] | 31 |
| Cuneus | [16, -80, 28] | 49 |
| Parietal_Sup | [-22, -70, 54] | 20 |

Region labels denote AAL.
